# CHK2 regulates MUS81-dependent DSBs in response to replication stress and BRCA2 deficiency

**DOI:** 10.1101/2022.10.08.511087

**Authors:** Eva Malacaria, Carolina Figlioli, Anita Palma, Sara Rinalducci, Maurizio Semproni, Annapaola Franchitto, Pietro Pichierri

## Abstract

MUS81 is a structure-specific endonuclease that processes DNA intermediates in mitosis and in S-phase following replication stress. MUS81 is crucial to cleave deprotected reversed forks in BRCA2-deficient cells. However, how MUS81 is regulated during replication stress in human cells remains unknown.

Our study reveals that CHK2 binds the MUS81-EME2 complex and positively regulates formation of DSBs upon replication stress or in the absence of BRCA2. The association with MUS81 occurs through the FHA domain of CHK2 and is disabled by the I157T mutation but not by the R117A mutation. The CHK2-MUS81 complex forms downstream fork reversal and degradation, and phosphorylation of MUS81 at CHK2-targeted sites is crucial to introduce DSBs at deprotected replication forks ensuring the replication fork recovery in BRCA2-deficient cells.

Collectively, our work sheds light into the regulation of the MUS81 complex and identifies a novel function of the ATM-CHK2 axis in the response to deprotected replication forks in the absence of BRCA2.

## INTRODUCTION

The replication stress response is critical for the maintenance of genome integrity and works as an anti-cancer barrier in human cells (1–4). One of the functions of this response is protection of the replication fork from extensive degradation and collapse (3, 5). To this end, perturbed replication forks are targeted by multiple proteins that coordinate a checkpoint response with fork remodelling and protection in response to sustained arrest (6–8). One of the critical steps towards fork protection is formation of a reversed fork (8). Replication fork reversal involves the action of multiple enzymes, including the key activity of SMARCAL1 and the homologous recombination factor RAD51 and BRCA2 (7). BRCA2 is crucial during homologous recombination to promote RAD51 nucleofilament assembly (9) and is mutated in the majority of familial breast cancers (10, 11). BRCA2 and RAD51 are also required to protect the reversed fork from unscheduled degradation by the MRE11/EXO1 nucleases (7, 8). Loss of BRCA2, indeed, impairs fork protection by RAD51 and stimulates a pathological degradation of nascent strands that eventually results in activation of a recombination-based replication recovery through break-induced replication pathways (12–14).

Pathological degradation and cleavage of reversed forks has been correlated with sensitivity of BRCA2-deficient cells to cisplatin or PARP inhibitors (15–17). Hence, understanding of this mechanism and its regulation has striking relevance also for cancer therapy. Unfortunately, little is known about the regulation of the pathological fork recovery pathway taking place in the absence of BRCA2.

In BRCA2-deficient cells, one of the critical steps of deprotected fork processing is the cleavage of a flap DNA intermediate by the MUS81-EME2 complex downstream MRE11/EXO1 activity (13, 18). The function of MUS81 in BRCA2-deficient cells is essential to replication fork recovery and to promote viability under conditions of normal or perturbed replication (13, 18).

MUS81 is a structure-specific endonuclease that cleaves recombination and replication DNA intermediates generating DSBs. MUS81 forms a heterodimeric complex with Eme1(Mms4) in yeast while human MUS81 can heterodimerize also with EME2 especially in S-phase and to process stalled forks (19). MUS81-dependent cleavage needs to be tightly regulated during cell cycle to avoid unscheduled DSBs formation in S-phase or insufficient resolution in G2/M (20). In human cells, the MUS81-EME1 complex activity in G2/M is regulated by CDK1, PLK1 and CK2 (20, 21). In contrast, little is known on how the MUS81 complex is regulated during replication, especially in response to replication stress and in pathological conditions, such as in BRCA2-deficient cells.

In fission yeast, Mus81/Eme1 function is controlled by Cdc2^CDK1^ and Chk1 kinases in response to DNA damage, while the checkpoint kinase Cds1^CHK2^ interacts through its FHA domain with MUS81 and negatively regulates its function during S-phase in S. pombe (22–25). In human cells, both CHK1 and CHK2 are activated during replication stress, however, CHK1 is the true functional Cds1 homologue although structurally CHK2 is the homologue of Cds1 (26).

Fork reversal and degradation generate free DNA ends that might activate CHK2 through ATM, in addition to ATR. However, whether CHK2 is involved in the pathological processing of perturbed forks upon replication stress and in the absence of BRCA2 is poorly investigated and is unknown if a functional MUS81-CHK2 crosstalk exists in human cells.

Therefore, in this study, we combined biochemical and single-cell assays to determine if CHK2 controls the formation of MUS81-dependent DSBs at replication forks and whether CHK2 and MUS81 functionally interact upon sustained arrest or loss of BRCA2.

Our study demonstrates that human CHK2 and MUS81 physically interact through the FHA domain, and that the I157T mutation significantly reduces this interaction, whereas the R117A mutation does not. Additionally, we show that CHK2 plays a role in regulating the formation of MUS81-dependent DSBs when replication forks are subjected to prolonged arrest or destabilization due to BRCA2 deficiency. Furthermore, we identify MUS81 residues phosphorylated by CHK2 that are functionally involved in DSBs formation at reversed replication forks in BRCA2-deficient cells. These findings reveal a critical regulatory mechanism that controls the MUS81 complex at demised replication forks, indicating that CHK2 activity and its binding with MUS81 govern the cleavage of reversed forks in cells lacking BRCA2.

## RESULTS

### MUS81-CHK2 interaction in response to replication stress involves the FHA domain of CHK2 and is disrupted by the I157T CHK2 mutation

To investigate the functional correlation between MUS81 and CHK2 in human cells, we first examined their possible association in a complex by performing immunoprecipitation experiments in HEK 293T cells transiently transfected with GFP-MUS81 and HA-CHK2 expression plasmids. Since MUS81 participates in the resolution of DNA intermediates in S-phase during replication stress and in mitosis, we treated cells with hydroxyurea (HU) or nocodazole (NOC) to accumulate cells in S-phase and mitosis, respectively (21). MUS81 was detected in the anti-HA immunoprecipitates from asynchronous and mitosis-enriched cells, however, HU increased MUS81-CHK2 complex formation 5-fold respect to untreated and 2-fold respect to NOC-treated cells (Fig. 1A). We then investigated the formation of the MUS81-CHK2 complex in shMUS81 MRC5 cells stably complemented with Flag-tagged wild-type MUS81 (MUS81^wt^) where the ectopic MUS81 is expressed at close-to-endogenous levels (21). CHK2 was detected in anti-Flag-MUS81 immunoprecipitates obtained from MRC5^wt^ cells while it was undetected in anti-Flag immunoprecipitation from the parental shMUS81 cells (Fig. 1B). The association between MUS81 and CHK2 was also confirmed at the single-cell level by proximity-ligation assay (PLA). As shown in Figure 1C, MUS81-CHK2 PLA-positive nuclei were detected in untreated MUS81^wt^ cells. Notably, replication stress increased both the number of PLA-positive nuclei and spots. In contrast, PLA signals were undetectable in shMUS81 cells (Fig. 1C).

**Figure 1.**
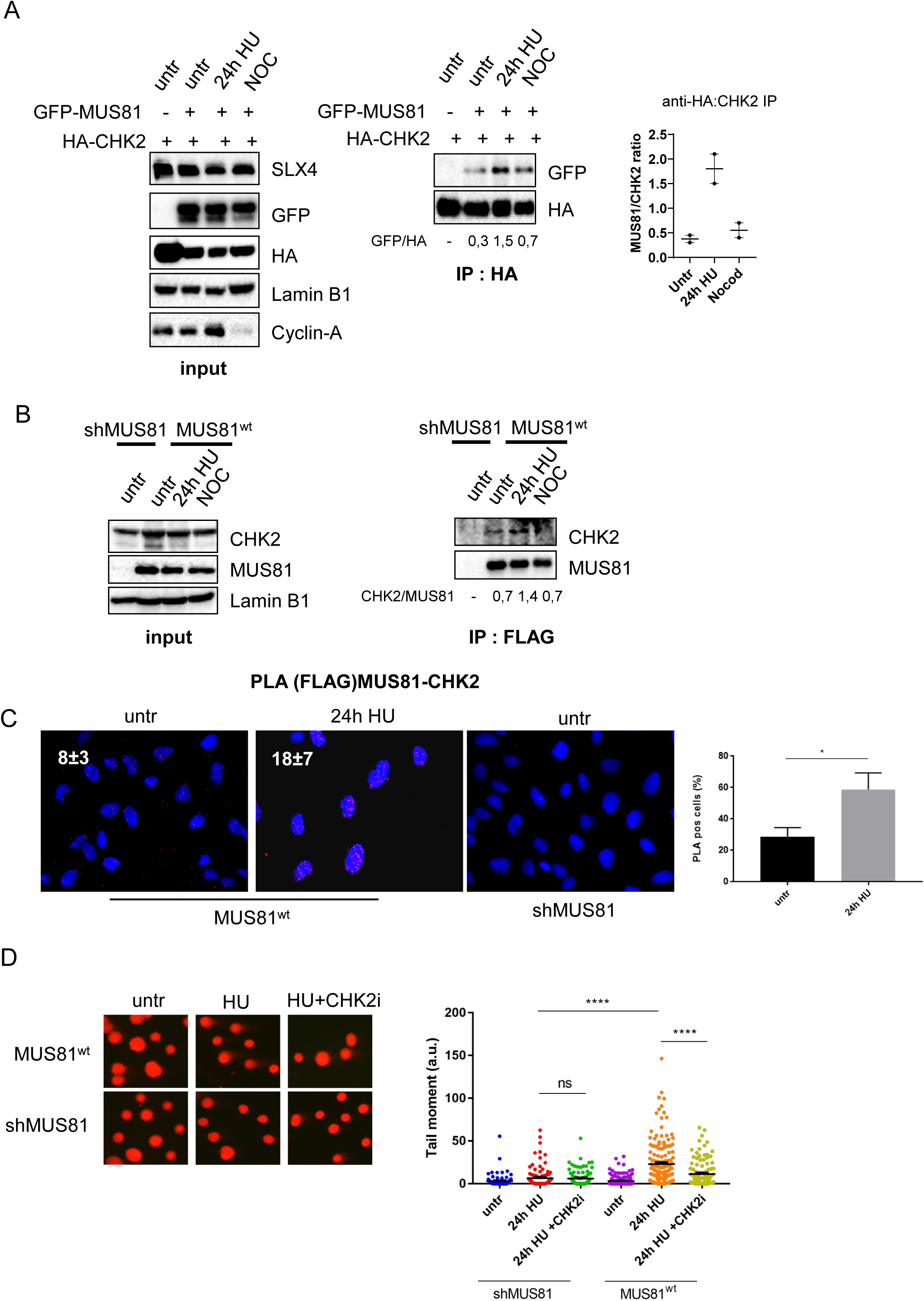
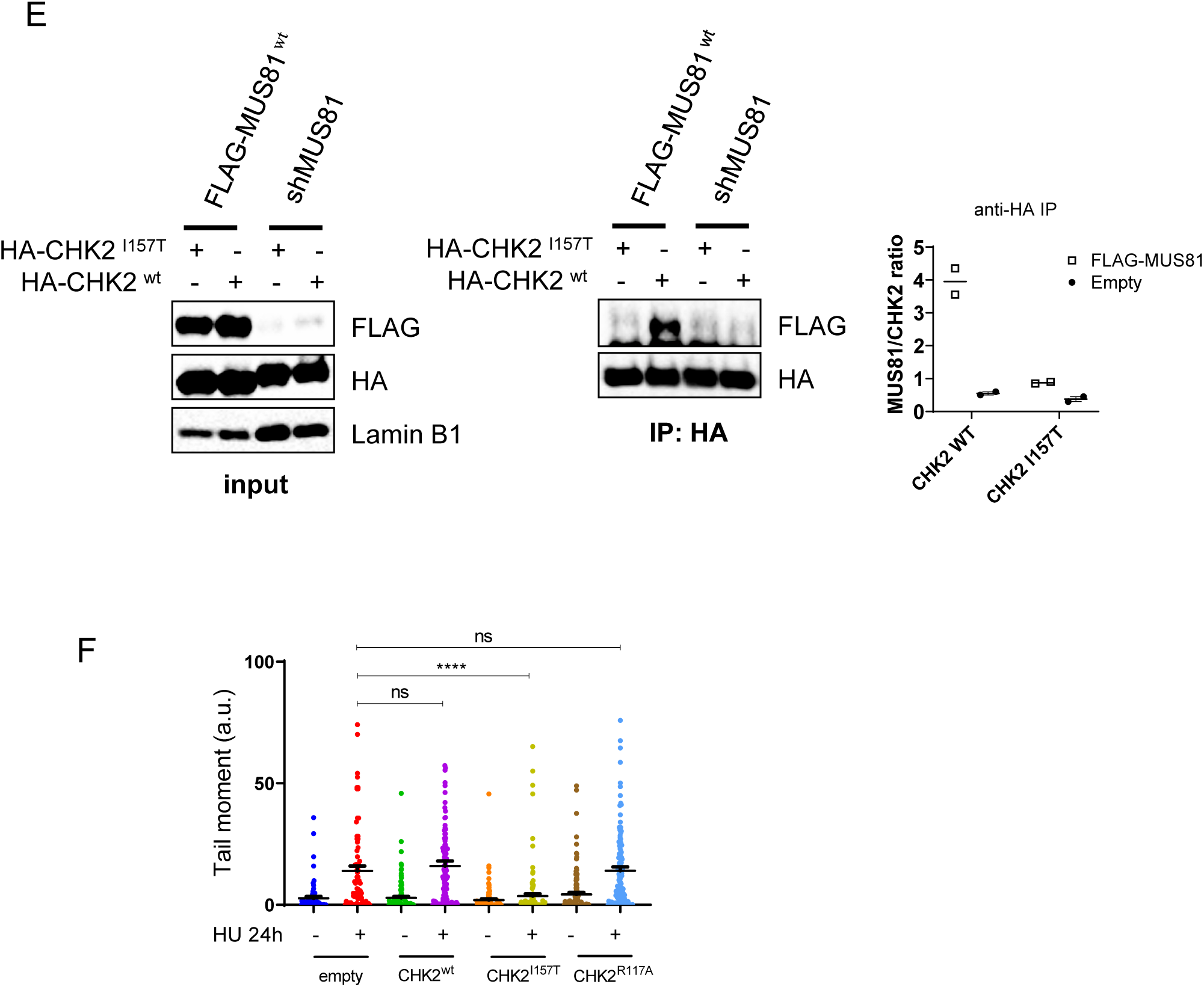
CHK2 form a complex with MUS81 that is stimulated upon replication stress and support MUS81-dependent DSB formation. A) HEK293T cells were transfected as indicated and 48h after treated with HU 2mM or nocodazole for 24h. After lysis, CHK2 or MUS81 were immunoprecipitated using anti-HA as indicated. The numbers below the IP represent quantification in arbitrary units of amount of MUS81 or CHK2 coimmunoprecipitating normalised on the fraction of CHK2 or MUS81 respectively. In the graph are presented mean from duplicated experiments. B). Endogenous MUS81 and CHK2 form a complex. MRC5 shMUS81 cells stably complemented with Flag-MUS81 were used to immunoprecipitate MUS81 in cells treated with HU or accumulated in mitosis with nocodazole. The numbers below the IP represent quantification in arbitrary units of amount of CHK2 coimmunoprecipitating normalised on the fraction of MUS81. C) MRC5 shMUS81 cells stably complemented with Flag-MUS81 were treated as indicated and analysed for the interaction between MUS81 and CHK2 using in situ PLA. Images show representative fields, and the inset shows the mean number of spots. The graph shows the number of PLA-positive nuclei (mean±SE; n=2. * = p < 0,5 ANOVA). D) MRC5 shMUS81 cells stably complemented or not with Flag-MUS81 were treated as indicated and subjected to neutral Comet assay to evaluate formation of MUS81-dependent DSBs. The panel shows representative images of comet tails for each condition. Quantification of Tail moment is reported in the graph. (ns = not significant; **** = p < 0,0001, Student’s t-test). E) MRC5 shMUS81 cells stably complemented with Flag-MUS81 were transfected with the indicated HA-CHK2 construct and analysed for the interaction between MUS81 and CHK2 by anti-HA co-immunoprecipitation. In the graph are presented mean from duplicated experiments. F) MRC5 shMUS81 cells stably complemented or not with Flag-MUS81 were transiently transfected with the indicated CHK2 construct, treated with HU, and subjected to neutral Comet assay to evaluate formation of MUS81-dependent DSBs. Quantification of Tail moment is reported in the graph. (ns = not significant; **** = p < 0,0001, Student’s t-test)

In human cells, MUS81 can heterodimerize with EME1 or EME2 (27). EME1 interacts with MUS81 throughout the cell cycle, while binding with EME2 is thought to occur predominantly during S phase (19). Despite a remarkable increase in the CHK2-MUS81 association after 24h of HU, both EME1 and EME2 were detected in the anti-CHK2 immunoprecipitates (Fig. S1A). To determine which of these two subunits contributes more to the interaction between MUS81 and CHK2 after replication stress, we performed RNAi-mediated knockdown of EME1 or EME2 and analysed the CHK2-MUS81 interaction using *in situ* PLA. Given that depletion of either EME1 or EME2 disrupts only one MUS81 complex, we expected that removing a key subunit might impair the interaction. However, depletion of EME1 did not affect the MUS81-CHK2 association, as shown by *in situ* PLA in HU-treated cells. In contrast, EME2 depletion led to a noticeable reduction in this interaction (Fig. S1B), suggesting that EME2 plays an important role in CHK2 association with MUS81 during replication stress.

Having demonstrated that CHK2 forms a complex with MUS81-EME2, we investigated whether this interaction was functional. Prolonged replication fork arrest by HU induces MUS81-dependent cleavage of stalled forks and unresolved replication intermediates (28–30). Since HU-induced replication stress enhanced MUS81-CHK2 interaction, we examined whether CHK2 kinase stimulates formation of MUS81-dependent DSBs. Using a neutral Comet assay, we assessed cells treated with HU for 24 hours, with or without the CHK2 inhibitor BML-277 (Fig. 1D). Cells expressing MUS81wt accumulated DSBs following HU treatment, and their levels were significantly reduced by CHK2 inhibition (Fig. 1D). In shMUS81 cells, few DSBs were detected after HU treatment, as expected, and CHK2 inhibition had no further effect (Fig. 1D).

In yeast, Cds1 interacts with Mus81 through its FHA1 protein-docking domain (23). Thus, we analysed whether, in human cells, the interaction of CHK2 with MUS81 after replication stress involved the FHA domain. To this end, we used a mutant form of CHK2 containing the I157T mutation in the FHA domain (31) This mutation, severely reduces protein interaction but not CHK2 activation (32). We performed immunoprecipitation assays in HEK 293T shMUS81 cells co-transfected with FLAG-MUS81 and HA-CHK2^wt^ or HA-CHK2^I157T^. After 24h of HU, MUS81 was readily detected in anti-HA immunoprecipitates from shMUS81 cells transfected with FLAG-MUS81 but not in the mock-transfected shMUS81 cells (Fig. 1E). However, co-immunoprecipitation between CHK2 and MUS81 was abrogated in the presence of the CHK2^I157T^ mutant. Consistent with CoIP data, we found that the wild-type CHK2-FHA domain fused to GST pulled down efficiently MUS81 from nuclear extracts, while the I157T mutation greatly decreased this binding (Fig. S2A). Functional integrity of the FHA domain of CHK2 has been shown to rely also on the R117 residue. Mutation R117A disrupts interaction with CHK2 partners, such as BRCA1 (33). Surprisingly, the FHA CHK2^R117A^ mutant, where the substitution of arginine 117 disrupts the phospho-dependent protein binding ability (32), did not affect interaction with MUS81 (Fig. S2B), suggesting that MUS81-CHK2 association does not necessarily involve prior phosphorylation as reported in yeast (ref 23).

Then, we investigated whether binding of the FHA-CHK2 domain with MUS81 was required to promote DNA breakage upon replication stress. To this aim, we performed neutral Comet assay in MRC5 cells transfected with HA-tagged CHK2^wt^, CHK2^I157T^ or CHK2^R117A^ (Fig. 1F). We reasoned that, since both mutants severely interfere with binding of CHK2 with multiple substrates but association with MUS81 is only abrogated by the I157T mutation, a specific defect in CHK2-MUS81 complex formation would have resulted in suppression of DSBs only in the CHK2^I157T^ mutant. Indeed, CHK2^I157T^ mutant overexpression abrogates DSBs accumulation after HU-induced replication stress while both CHK2^wt^ or CHK2^R117A^ overexpression failed to affect DSBs formation as compared with control transfected cells (empty) (Fig. 1F).

Together, these results demonstrate that MUS81 and CHK2 form a complex in response to replication stress-induced fork collapse. The CHK2-MUS81 interaction depends on the FHA domain of CHK2, which is disrupted by the I157T mutation but not by the R117A mutation. Additionally, CHK2 inhibition or interference with MUS81 binding to the CHK2 FHA domain prevents DSB formation under replication stress.

### Formation of the MUS81-CHK2 complex and activity of CHK2 promotes DSBs at deprotected forks

The MUS81 complex targets replication forks after their prolonged stalling but also cleaves deprotected reversed forks in the absence of BRCA2 or other replication caretakers (12, 13, 29, 34–36). We therefore investigated if interaction between CHK2 and MUS81 was involved in the generation of DSBs occurring in BRCA2-depleted cells. CHK2 inhibition or CHK2^I157T^ overexpression were used to test the effect of CHK2 activity and MUS81/CHK2 binding on DSBs formation respectively (Fig. 2A, B). As expected, treatment with HU for 2h induced a significant amount of DSBs in absence of BRCA2, as shown by neutral Comet assay. Exposure of cells to the BML-277 CHK2i during the HU treatment greatly reduced formation of DSBs. Similarly, overexpression of the FHA I157T CHK2 mutant greatly reduced the formation of MUS81-dependent DSBs occurring in BRCA2-depleted cells after the short HU treatment. Of note, in BRCA2-depleted cells, expression of the CHK2^I157T^ mutant or the absence of MUS81 each reduced DSBs although expression of the CHK2^I157T^ mutant appeared slightly more efficient in preventing DSBs (Fig S3). Consistent with a role of CHK2 in modulating the function of the MUS81 complex at deprotected forks, its activatory phosphorylation at T68 and that of ATM at S1981 were both elevated in the absence of BRCA2, especially at 2h of treatment (Fig. S4A). To determine if increased activation of ATM as seen in the absence of BRAC2 originated from stalled replication forks, we analysed recruitment of active ATM at nascent DNA by SIRF using an anti-pS1981ATM antibody (Fig. 2C). We expected more active ATM recruited to replication forks in absence of BRCA2 early after HU treatment. Notably, active ATM was recruited early at replication forks in BRCA2-proficient cells. Its association with these forks increased after 4 hours of HU treatment and subsequently declined. However, an increased amount of pS1981 ATM was observed at stalled replication forks in the absence of BRCA2 by SIRF (Fig. 2C). Furthermore, the proportion of fork-associated pS1981 ATM remained consistent even at subsequent time points. The analysis of pS1981ATM foci in chromatin fractionated cells following MUS81 depletion indicated that the elevated ATM activation occurs independently of the MUS81-dependent formation of DSBs in cells depleted of BRCA2 (Fig. S4B). In BRCA2-deficient cells, MUS81 activity is required for regressed fork cleavage and subsequent replication restart (13). Thus, we analysed whether inhibition of CHK2 or mutation of its FHA domain to abrogate the interaction with MUS81 could affect recovery from replication arrest in BRCA2-depleted cells. To this end, we treated BRCA2-depleted cells with CHK2i or transfected them with the I157T or R117A FHA-CHK2 mutant, which disables or not the CHK2-MUS81 interaction respectively (Fig. 2D). Replication fork recovery was evaluated as the residual number of stalled forks on stretched DNA fibres prepared from cells pulse labelled with CldU and IdU (Fig. 2D). Consistent with previous data (13), replication fork recovery was reduced in cells depleted of BRCA2 and was largely dependent on MUS81. Inhibition of CHK2 in BRCA2-depleted cells also increased the number of stalled forks after recovery from HU treatment and, notably, phenocopied MUS81 depletion (Fig. 2D). Interestingly, the residual number of stalled forks was similarly increased by expression of the I157T FHA-CHK2 mutant, which compromised MUS81 binding, while was not affected by the R117A FHA-CHK2 mutant that retains normal MUS81 binding.

**Figure 2.**
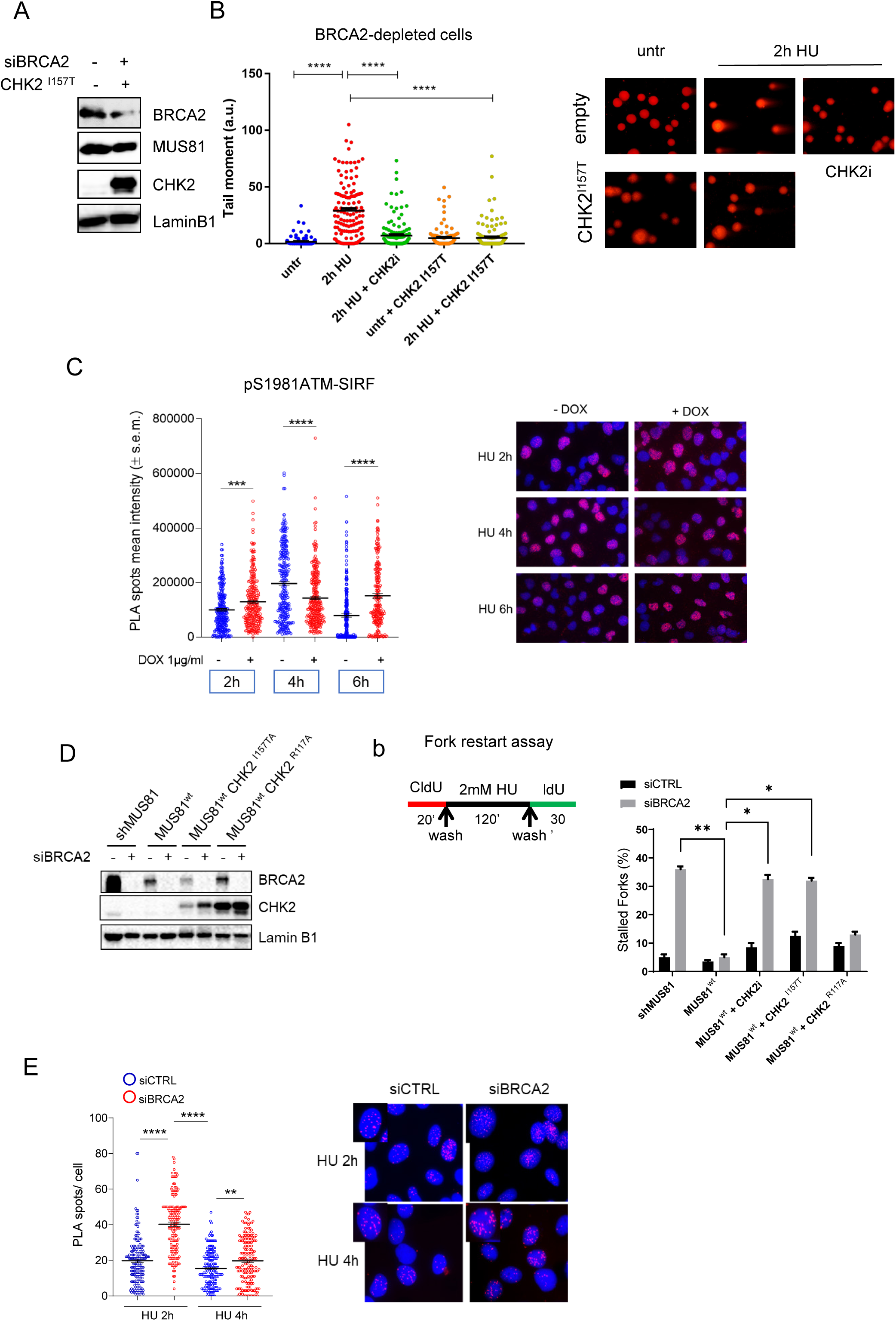
Activity of CHK2 and its interaction with MUS81 drives formation of DSBs at stalled forks in BRCA2-deficient cells. A) WB showing depletion of BRCA2 and expression of I157T-mutated CHK2 in MRC5 shMUS81 cells stably complemented with Flag-MUS81. B) MRC5 shMUS81 cells stably complemented with Flag-MUS81 were transfected with the indicated HA-CHK2 construct or treated with the BML-277 CHK2 inhibitor (CHK2i) prior to be exposed to HU 2mM and analysed for the presence of MUS81-dependent DSBs by neutral Comet assay. (**** = p<0.001, Student’s t-test). Representative images are shown. C) Analysis of fork recruitment of active ATM in BRCA2-depleted cells. Doxycycline-inducible shBRCA2 MRC5 cells were depleted of BRCA2 by treatment with dox, as indicated, and treated with HU for different time-points. Recruitment of active ATM at nascent DNA at fork was analysed by SIRF using anti-pS1981ATM antibody. (*** = p<0.01; **** = p<0.001, ANOVA). Representative of PLA images are shown. D) WB showing depletion of BRCA2 and expression levels of mutant CHK2 in MRC5 shMUS81 cells stably complemented with Flag-MUS81. Analysis of replication fork restart using DNA fibre assay. Cells were treated as indicated in the scheme on the top. The graph shown the percentage of stalled forks for each condition. (** = P < 0.01; *** = p<0.001, ANOVA test). E) Cells were transfected with scrambled siRNA (siCTRL) or siRNA directed against BRCA2 alone or in combination with siRNA against SMARCAL1 and treated with HU prior to the evaluation of MUS81-CHK2 interaction in situ by PLA. The graphs show the number of PLA spots for each condition (n=2, * = P < 0.5; **** = P < 0.0001; Mann–Whitney test). Representative images are shown.

We therefore concluded that activation of CHK2 and its interaction with MUS81 are required for the early DSBs formation occurring in BRCA2-depleted cells and to subsequently promote fork recovery. Thus, we wanted to determine if the interaction between CHK2 and MUS81 was stimulated by the absence of BRCA2. Hence, we performed PLA experiments in cells exposed to HU for different time-points. Interestingly, a significant CHK2-MUS81 interaction was observed 2 hours after replication arrest even in BRCA2-proficient (siCTRL) cells, and this interaction persisted at 4 hours (Fig. 2E). This finding may indicate that CHK2 and MUS81 form a complex even when MUS81 is not required to target perturbed replication forks. However, cells depleted of BRCA2 (siBRCA2) showed a significant increase in the number of CHK2-MUS81 PLA spots at 2h of HU that declined to wild-type levels at 4h (Fig. 3A). Combined PLA and EdU detection confirmed that most PLA spots occurred in S-phase in BRCA2-deficient cells, consistent with the CHK2-MUS81 interaction taking place during replication arrest (Fig. S5).

**Figure 3.**
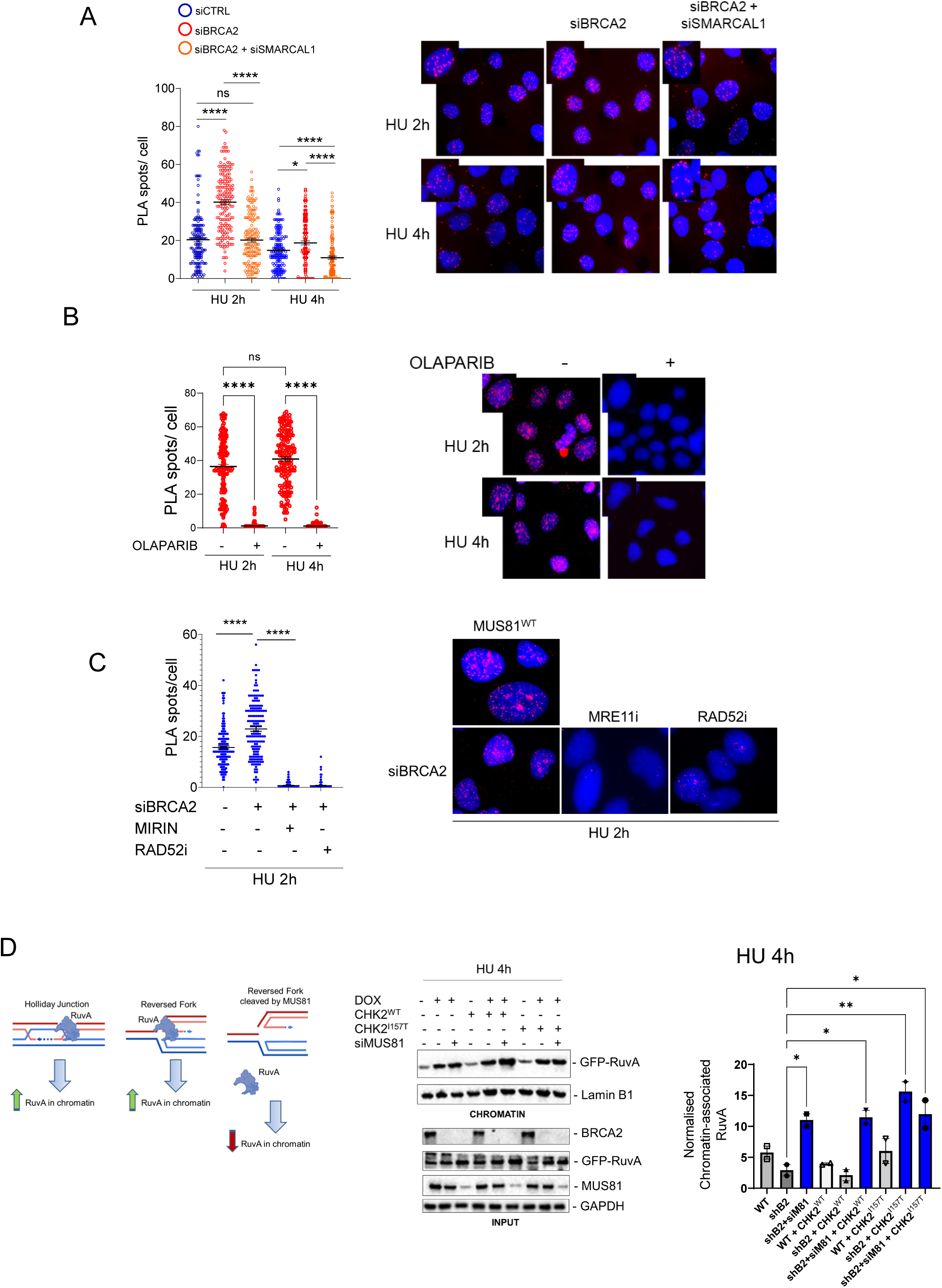
Interaction between MUS81 and CHK2 is dependent on fork reversal and processing. A) Cells were transfected with scrambled siRNA (siCTRL) or siRNA directed against BRCA2 alone or in combination with siRNA against SMARCAL1 and treated with HU prior to the evaluation of MUS81-CHK2 interaction in situ by PLA. The graphs show the number of PLA spots for each condition (n=2, * = P < 0.5; **** = P < 0.0001; Mann–Whitney test). Representative images are shown. C) Cells were depleted of BRCA2 by transfection with siRNAs and pre-treated with Olaparib before being challenged with HU as indicated. Interation between MUS81 and CHK2 was evaluated by in situ PLA. The graph shows the number of PLA spotes for each condition (n=2, **** = P < 0.0001; Mann–Whitney test). Representative images are shown. D) Cells were transfected with scrambled siRNA (siCTRL) or siRNA directed against BRCA2. Cell lysates were subjected to immunoblotting with indicated antibodies. Lamin B1 was used as loading control. Single-cell analysis of CHK2-MUS81 interaction by in situ PLA. Cells were treated as indicated in the graph. Graph shows the number of PLA spots per cell. (n=2, **** = P < 0.0001; Mann– Whitney test). Representative images are shown. D) Cartoon exemplifying the use of RuvA as proxy for the presence of 4-way junctions and expected results if MUS81 is cleaving or not reversed forks. Dox-regulated shBRCA2 MRC5 cells were transfected with plasmids expressing GFP-RuvA and the indicated HA-CHK2 construct. After 48h, cells were treated with HU and analysed for the presence of RuvA in chromatin by WB of chromatin fractions with the indicated antibodies. RuvA was detected using anti-GFP while CHK2 using anti-HA antibodies. Input shows expression levels of RuvA and CHK2 or MUS81 depletion. The graph shows the normalized amount of RuvA in chromatin for each condition. In blue, are highlighted conditions in which MUS81 is not working or absent. Values are presented as means ± S.E. (n=2, not significant pairs are not labelled; *=p<0.5; **=p<0.1; ANOVA)

These results indicate that the ATM-CHK2 axis is activated to promote MUS81-dependent DSBs and that formation of the CHK2-MUS81 complex is essential for correct function of the MUS81 complex in BRCA2-depleted cells.

### The interaction between CHK2 and MUS81 takes place after fork reversal to target deprotected replication forks in absence of BRCA2

In BRCA2-deficient cells, the MUS81 complex targets deprotected forks after their reversal (13). To determine whether, in BRCA2-deficient cells, formation of the CHK2-MUS81 complex occurred downstream fork reversal, we performed CHK2/MUS81 PLA after transient depletion of SMARCAL1, one of the crucial enzymes involved in this pathway (37).

As anticipated, transfection with siBRCA2 oligos resulted in increased formation of a CHK2-MUS81 complex early after HU treatment compared to cells transfected with siCTRL oligos (Fig. 3A). Concomitant depletion of SMARCAL1 and BRCA2 greatly reduced the interaction of MUS81 with CHK2 at all time-points (Fig. 3B). However, depletion of SMARCAL1 did not suppress interaction between MUS81 and CHK2 completely. Thus, we performed PLA experiments in cells pre-treated with Olaparib, a condition that prevents fork reversal (38). Of note, Olaparib pre-treatment completely suppressed the formation of the CHK2-MUS81 complex as detected by PLA in BRCA2-depleted cells at 2 and 4 hours of HU treatment (Fig. 3C). In BRCA2-depleted cells, MUS81 acts downstream the pathological degradation of the reversed forks by MRE11/EXO1, and requires the presence of RAD52 (13, 14, 39, 40). Thus, to further demonstrate that the CHK2-MUS81 complex in engaged post fork reversal and degradation, we performed PLA experiments in BRCA2-depleted cells in which MRE11 was inhibited with Mirin or RAD52 with EGC (RAD52i; (39)). Again, multiple PLA spots were observed in BRCA2-depleted cells after HU treatment indicative of the CHK2-MUS81 interaction, and, interestingly, either Mirin or the RAD52i completely suppressed formation of PLA spots (Fig. 3B).

In the absence of a functional MUS81 complex, the number of resected, but unbroken, reversed forks increases in BRCA2-deficient cells, and this correlates to reduced replication restart (13); a phenotype that we observe when the interaction with CHK2 is disrupted by the I157A mutation or CHK2 is inhibited (see Fig. 2D). We previously showed that ectopically expressed bacterial RuvA, an avid binder of HJs, accumulates in chromatin under conditions stimulating formation of reversed forks (40). Therefore, we used chromatin recruitment of RuvA as an indicator of the presence of reversed forks in cells that stably express a doxycycline-regulated shBRCA2 cassette. These cells were transiently transfected with the CHK2 mutants, either alone or in combination with siRNA targeting MUS81, before being treated with HU to induce replication fork stalling (Fig. 3D).

As shown in Figure 3D, HU treatment induced the presence of chromatin-associated RuvA in BRCA2-proficient cells (WT cells; no DOX), consistent with the presence of HJ-like structures, including reversed forks. Depletion of BRCA2 by DOX, slightly reduced the presence of RuvA in chromatin, which was increased by concomitant depletion of MUS81, as expected if chromatin-bound RuvA marks reversed forks that persist if left uncleaved (Fig. 3D). Interestingly, in BRCA2-depleted cells, expression of the CHK2^I157T^ FHA mutant, which does not bind MUS81, also greatly increased the amount of RuvA in chromatin but not if combined with MUS81 depletion.

Nascent ssDNA is formed as the result of resected reversed forks (13, 40–42). CHK2 inhibition or inability to interact with MUS81 did not significantly decrease the amount of exposed nascent ssDNA (Fig S6) in agreement with the function of MUS81 downstream fork degradation by MRE11-EXO1 (13).

These results indicate that, in BRCA2-depleted cells, the formation of the MUS81-CHK2 complex occurs downstream fork reversal and after fork degradation, possibly stimulating correct targeting of reversed forks downstream fork degradation.

### CHK2-dependent phosphorylation of MUS81 at S97 controls formation of DSBs under replication stress

FHA domain mediates the recognition of CHK2 substrates (43). Thus, we wondered if MUS81 could be a CHK2 target also in human cells. To test if MUS81 is a CHK2 substrate, we performed an *in vitro* phosphorylation assay. As substrate we used the 76-206 N-terminal region of human MUS81 that contains an essentially unstructured region including SLX4 binding residues (44, 45) and several CHK2 putative phosphorylation sites (21). The purified N-terminal fragment of MUS81 fused to GST was incubated with recombinant CHK2 kinase, subjected to SDS-PAGE and analysed by MS/MS (Fig. 4A). Mass spectrometry identified two major phosphorylation sites, Thr 86 and Ser 97, in the MUS81 fragment *in vitro* (Table 1 and Figures S7 and S8). To determine if MUS81 phosphorylation at these residues had any functional relevance in response to replication stress, we performed neutral Comet assay in shMUS81 cells transiently transfected with wild-type MUS81 or its unphosphorylable (T86A, S97A) or phosphomimetic (T86D, S97D) mutants (Fig. 4B). Notably, the presumed CHK2-unphosphorylable T86A and S97A MUS81 mutants were unable to facilitate HU-induced accumulation of DSBs. Additionally, combined CHK2 inhibition did not further decrease DSB levels, indicating that phosphorylation at these sites is CHK2-dependent within the cell. The phosphomimetic T86D MUS81 mutant showed a wild-type ability to induce DSBs after prolonged HU treatment but not in untreated conditions, while the S97D MUS81 mutant caused a striking amount of DSBs formation already under unchallenged replication.

**Figure 4.**
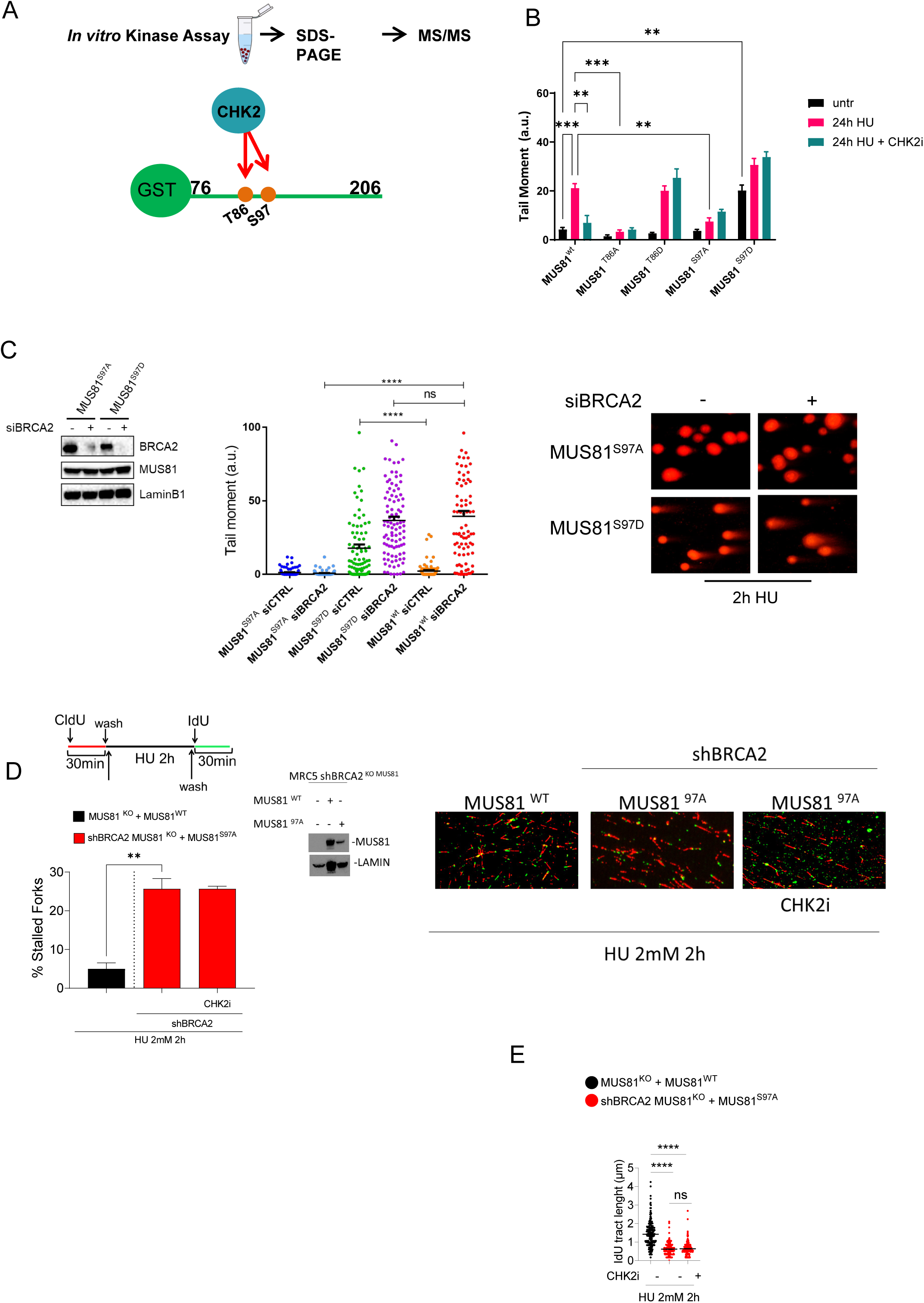
Abrogation of phosphorylation of MUS81 at CHK2 sites impairs the function of MUS81 after replication stress. A) Scheme outlining the identification of *in vitro* MUS81 phosphosites. GST-fused N-terminal MUS81 fragment was incubated with recombinant CHK2 and subjected to MS/MS analyses to identify phosphopeptides and residues. B) MRC5 shMUS81 cells were transfected with plasmids expressing different MUS81 phosphomutant for each CHK2 site identified by MS. After 48h, cells were treated as indicated. Formation of DSBs was analysed by neutral Comet assay. The graph shows the mean Tail moment ± S.E. for each condition. (ns = not significant; *** = p<0.01; **** = p<0.001 Student’s t-test). C) WB showing BRCA2 levels in MRC5 shMUS81 cells transiently transfected with the indicated MUS81 mutant and BRCA2 siRNAs. The graph shows the mean Tail moment ± S.E. for each condition. (ns = not significant; **** = p<0.001 Student’s t-test). Representative images are shown. D) Analysis of replication fork restart using DNA fibre assay. Cells were treated as indicated in the scheme on the top. The graph shown the percentage of stalled forks for each condition. Inducible shBRCA2 MUS81 knockout MRC5cells were transfected with MUS81 WT or MUS81 97A. Cell lysates were subjected to immunoblotting with indicated antibodies. Lamin B1 (LAMIN) was used as loading control. Representative images of DNA fibres fields are presented. (**P < 0.01; ANOVA). E) Analysis of IdU tract length of restarted forks. Length of the green tracks was measured in 200 individual forks. Mean values ± SE are represented as horizontal black lines. (ns = not significant; **** P < 0.0001; Mann–Whitney test)

**Table 1.** Phosphorylation sites identified in the N-terminal MUS81 region by in vitro CHK2 kinase assay.

| Residue position | Residue type | Phosphopeptide Sequence | Mascot score | m/z (charge) | Fragmentation type | Retention time |
| --- | --- | --- | --- | --- | --- | --- |
| 97 | Ser | TSGGDHAPDSP**S**pGENSPAPQGR | 107 | 734.31 (3+) | ETD | 14.4 min |
| 86 | Thr | HRT**p**SGGDHAPDSPSGENSPAPQGR | 52 | 832.04 (3+) | CID | 13.9 min |

We next focused on S97 because the phenotype of the phosphomimetic S97D mutation indicated that it could be involved in replication processes that require strict regulation. Therefore, we examined whether MUS81-S97A and S97D could affect DSBs formation at deprotected forks by performing a neutral Comet assay upon depletion of BRCA2 and transient transfection of the corresponding unphosphorylable or phosphomimetic MUS81 mutants (Fig. 4C). We did not detect more DSBs in MUS81^S97A^ siBRCA2-transfected cells if compared with cells transfected with control siRNA (Fig. 4C), suggesting that S97 needs to be phosphorylated in the absence of BRCA2. However, MUS81-S97D expression significantly increased DSBs in BRCA2-proficient cells after a 2-hour HU treatment. Interestingly, in BRCA2-depleted cells, MUS81-S97D did not further increase DNA breakage compared to BRCA2-proficient cells, suggesting that maintaining high phosphorylation on this residue is necessary under these conditions. As reported consistently in previous reports (13, 29), a short treatment with HU failed to induce any detectable DSBs in wild-type cells (Fig. 4C).

Neutral Comet assay suggests that S97 of MUS81 needs to be phosphorylated by CHK2 in BRCA2-depleted cells. Since, in these conditions, DSBs are required for the recovery of stalled replication forks, we quantified the percentage of replication forks stalling by DNA fiber assay after HU treatment. To this end, we used MUS81^KO^ cells stably-expressing a dox-regulated shBRCA2 cassette and transfected transiently with the S97 CHK2 mutants (shBRCA2 MUS81^KO^ ((46); Fig 4D)

Although the number of stalled forks is low after 2h of HU, the fraction of stalled forks in BRCA2-depleted cells increased significantly after expression of MUS81^S97A^, phenocopying what was observed after CHK2 inhibition (see Fig 2D). Interestingly, inhibition of CHK2 in BRCA2-depleted cells expressing MUS81^S97A^ failed to further increase the number of stalled forks (Fig. 4D). In addition to affecting the number of stalled forks that restart DNA synthesis, expression of MUS81-S97A in BRCA2-depleted cells also reduced the efficiency of fork restart, leading to shorter IdU tracts (Fig. 4E). These results indicate that CHK2 can target MUS81 at multiple residues and that phosphorylation status of these residues is critical to regulate the formation of MUS81-dependent DSBs and to promote restart of stalled replication forks in BRCA2-depleted cells.

### CHK2-MUS81 interaction promotes survival of BRCA2-deficient cells upon replication stress

MUS81 cleaves intact forks and partially regressed forks in wild-type cells or BRCA2-depleted cells, respectively (13, 28), leading to rescue after genotoxic stress or replication fork deprotection. To test whether loss of CHK2-MUS81 complex formation was required for survival of BRCA2-deficient cells after HU treatment, we quantified the viability of cells after transient expression of the wild-type form of CHK2 or the CHK2^I157T^ mutant disabling the interaction of the FHA domain with MUS81 using a short-term viability assay. We used MUS81^KO^ cells stably-expressing a dox-regulated shBRCA2 cassette (in brief, MUS81^KO^) or their non-edited counterpart expressing wild-type MUS81 (46). As reported by others (13), concomitant loss of BRCA2 and MUS81 sensitized cells to HU treatment (Fig. 5A). Expression of the CHK2^I157T^ mutant increased HU sensitivity in BRCA2-depleted cells also in the presence of the wild-type form of MUS81 as compared with cells expressing CHK2^wt^ (Fig. 5A). Interestingly, expression of the CHK2^I157T^ mutant did not strikingly enhance sensitivity to HU in MUS81^KO^, BRCA2-depleted, cells (Fig. 5A). Similarly, expression of the MUS81-S97A mutant in MUS81^KO^, BRCA2-depleted cells, failed to revert HU sensitivity. Thus, either MUS81 KO, abrogation of the interaction with CHK2 or of MUS81 phosphorylation is sufficient to induce a similar hypersensitivity to HU in BRCA2-depleted cells.

**Figure 5.**
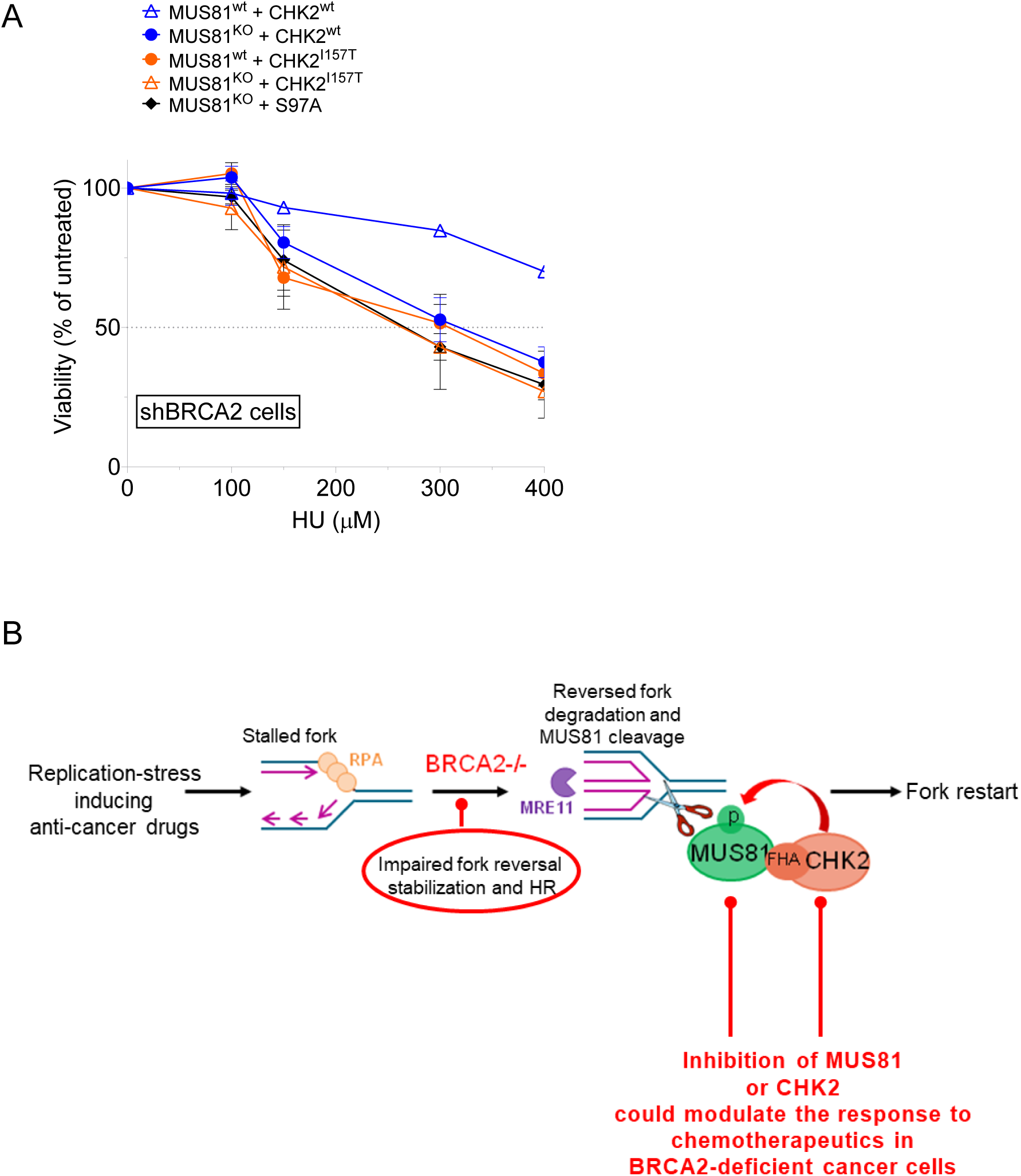
The absence of MUS81 or inability to form the CHK2-MUS81 complex similarly affects HU sensitivity of BRCA2-depleted cells. A) The wild-type or FHA mutant form of CHK2 was transiently expressed in BRCA2-depleted MRC5 cells wild-type or KO for MUS81. Viability was assessed by MTT assay after treatment with HU for 4 days. Each point represents the average number of counts from four independent experiments. B) Model of CHK2-MUS81 complex function in normal or BRCA2-depleted cells. See discussion for details.

## DISCUSSION

In this work, we have shown that activation of the MUS81 complex during prolonged replication fork arrest or pathological responses to fork stalling requires activity of CHK2 and its association with MUS81 through the FHA domain. CHK2 activity, binding to and phosphorylation of MUS81 are critical events for the processing of reversed forks into DSBs when BRCA2 is absent, contributing to replication fork recovery and viability (Fig. 5B). This defines a previously unappreciated regulatory mechanism of the MUS81 complex during replication fork cleavage and restart upon replication stress and deprotection of reversed forks.

Since formation of DSBs at the replication fork can lead to gross chromosomal rearrangements, activity of the MUS81 endonuclease is normally blocked or negatively-regulated in S-phase (27). In yeast, the Mus81 complex is inhibited during normal replication or following transient arrest by Cds1, while checkpoint kinases stimulate its function, cooperating with Cdks, in response to DNA damage in S-phase or, in budding yeast, when Rqh1 is absent (22, 24, 25, 47, 48).

In human cells, phosphorylation of EME1 by CDK1/PLK1 and of MUS81 by CK2 on Serine 87 restrict the activation of SLX4-MUS81-EME1 complex in late G2/M, preventing aberrant cleavage in S-phase (21, 49). Here, we find that HU-induced replication stress leads to the formation of a CHK2-MUS81 complex. Notably, MUS81-dependent DSBs arising at perturbed replication forks are EME2-dependent, suggesting that CHK2 may work to regulate preferentially MUS81 when complexed with EME2, consistent with previous work showing EME2 relevance in S-phase (19, 30, 50). However, previous data from our group and others implicated MUS81/EME1 in the response to CHK1 inhibition or oncogene-induced replication stress (34–36) indicating that different and specific complexes of MUS81 might form also in S-phase under pathological conditions.

In yeast, the inhibitory interaction between Cds1 and MUS81 involves the FHA domain of Cds1 (23, 24). Our data clearly indicate that the CHK2-FHA domain is responsible for MUS81 binding also in humans, although we find that CHK2 regulates positively MUS81-dependent DSBs formation after replication stress. Although binding of Cds1 to Mus81 is phosphodependent (24), we failed to identify a phosphorylation-dependent CHK2-FHA-MUS81 interaction. However, FHA domains show versatile and broad specificity, binding canonical phosphopeptides as well as non-canonical and non-phosphorylated sequences (51, 52), possibly suggesting that the CHK2-MUS81 association involves a FHA-mediated but phospho-independent interaction occurring in a distinct FHA peptide-binding surface (32, 43). Interestingly, we find that the FHA ^I157T^ mutation largely impairs MUS81-CHK2 interaction and MUS81-dependent DSBs under replication stress. In contrast, but consistent with a phospho-independent interaction, the FHA^R117A^ mutant, which impairs the affinity for target phosphates and abolishes CHK2 interaction with BRCA1 (32), does not affect MUS81 binding and function in response to replication stress, arguing that the phenotype observed in cells overexpressing the FHA mutant conferring a defective interaction with MUS81 might be specific.

The MUS81/EME2 complex is crucial also to overcome fork deprotection induced by loss of BRCA2 (13). However, it is also crucial that MUS81/EME2 targets only degraded replication forks. Our findings reveal that CHK2-dependent regulation of MUS81 is required for DSBs formation and fork restart in BRCA2-deficient cells. Interestingly, association between CHK2 and MUS81 is stimulated in BRCA2-deficient cells and CHK2 or ATM activatory phosphorylation at T68 or S1981, respectively, are elevated in the absence of BRCA2 early after replication arrest. Moreover, phosphorylated ATM can be detected at stalled forks independently of the presence of MUS81. These results suggest that ATM-CHK2 is activated prior to DSBs formation in BRCA2-deficient cells, consistent with an upstream role in regulating MUS81/EME2 function. Interestingly, active, S1981-phosphorylated, ATM can be detected at stalled replication forks even in BRCA2-proficient cells where DSBs are undetectable. This last observation is consistent with the recognition of DNA ends of reversed forks by DSB binding factor, like KU (53). Indeed, MUS81-CHK2 PLA spots can be detected also in BRCA2-proficient cells. This indicates that the MUS81-CHK2 complex is assembled whenever forks get stalled but is kept inactive or that a fraction of stalled forks are unstable even in the presence of BRCA2, stimulating assembly of the complex. Further studies will be required to discriminate between these two hypotheses. Formation of the MUS81-CHK2 interaction is greatly reduced when fork reversal is impaired. Consistent with this, preventing MRE11 from binding to or degrading the reversed fork interferes with MUS81-CHK2 interaction in the absence of BRCA2.

Upon loss of BRCA2, MUS81 acts downstream CtIP, MRE11 and EXO1 nucleases to mediate cleavage of a flapped DNA at the reversed forks (13). Thus, our data indicate that the MUS81-CHK2 complex is assembled downstream fork remodelling to ensure that cleavage occurs only at specific DNA intermediates.

In absence of MUS81-dependent cleavage, extensively resected forks cannot properly restart (13). Consistent with this, loss of CHK2-dependent MUS81 regulation conferred by the FHA I157T mutation impairs cleavage of stalled replication forks with their significant accumulation and severe consequences on the ability to restart replication and overcome replication stress.

Mutations of ATM or CHK2 predispose to breast cancer and the ATM-CHK2 pathway has been shown to support MUS81 activity to generate DSBs in response to cisplatin treatment in a human breast cancer model (54). Although, the missense CHK2 I157T mutation previously associated to Li-Fraumeni syndrome (31, 55), has been now assigned as a SNP, it is linked with breast cancer (odds ratio = 1.4) and increases the risk of breast cancer if carriers already have deleterious mutations of CHK2 and BRCA1/2 (56). Our data demonstrate that this variant specifically abrogates CHK2 binding to MUS81 and impairs fork recovery, consistent with previous studies showing defective binding to other checkpoint regulators such as BRCA1. Thus, this allele serves as a valuable tool to functionally interrogate CHK2’s role in regulating endonuclease activity during replication stress, independently of its clinical penetrance.

Of note, CHK2 can target at least two distinct MUS81 residues, *in vitro*. Mutations abrogating phosphorylation at these two residues suppress MUS81-dependent DSBs after prolonged replication arrest in wild-type cells suggesting that modification of these sites is functional. Interestingly, the two putative CHK2 phosphorylation sites localise close to the N-terminal region of MUS81 involved in the interaction with SLX4 that acts to relieve the auto-inhibitory binding between the Hairpin-helix-hairpin domain of MUS81 and the EME1 subunit (45). Given the ample structural similarities between EME1 and 2 (57), it is tempting to speculate that phosphorylation by CHK2 might promote a conformational change similar to that induced by interaction with SLX4 thus leading to an active MUS81-EME2 complex. Inability of S>A MUS81 mutants to induce DSBs and elevated DSBs in the S97D mutant, even in untreated conditions, may be consistent with the presence of an inactive CHK2-MUS81 complex requiring regulated phosphorylation for its full activation.

In conclusion, this work sheds light on regulation of the MUS81/EME2 complex and identifies how CHK2 regulates the response of BRCA2-deficient cells to deprotected forks. Since activation of MUS81 at deprotected forks has been correlated with response to Olaparib (15, 16) and inactivating CHK2 mutations can be found in breast cancer (58, 59), our work might be of value for the identification of novel biomarkers of PARPi resistance.

In addition, because CHK2 inhibitors are being evaluated in preclinical studies, our mechanistic evidence may be helpful for an educated combination of DDR inhibitors to avoid “self-inactivating” combinations of inhibitors.

## ACKNOWLEDGMENTS

We are grateful to Drs Domenico Delia and Giacomo Buscemi for the gift of the pcDNA3-HA-Chk2 plasmid. We are grateful to Prof. Massimo Lopes for scientific discussion. This work was supported by the Italian Ministry of Health “Ricerca Finalizzata” to P.P. (grant n. RF-2016-02362022) and by investigator grants from Associazione Italiana per la Ricerca sul Cancro (AIRC) to P.P. (IG n. 21428) and to A.F. (IG n. 19971).

## AUTHOR CONTRIBUTIONS

A.P. performed all the biochemical experiments and functional characterisation of the MUS81-CHK2 complex formation. E.M. and C.F. performed the analysis of fork recruitment in vivo, analysis of ATM-CHK2 activation, analysis of RuvA chromatin recruitment, DNA fibers assays and most of the functional analysis of the MUS81 phosphorylation mutants. M.S. assisted with the purification of protein fragment in bacteria and with CoIP. S.R. performed the analysis of MUS81 phosphorylation and MS to identify phosphorylated residues. A.P., E.M., S.R., analysed data and contributed to designing the experiments and writing the paper. A.F. and P.P designed experiments, analysed data, supervised research and wrote the paper. All authors approved the paper. E.M. and A.P. contributed equally to this work and should be regarded as joint first authors.

## MATERIALS AND METHODS

### Cell lines and culture conditions

The MRC5SV40 cells were maintained in Dulbecco’s modified Eagle’s medium (DMEM; Life Technologies) supplemented with 10% foetal bovine serum (Boehringer Mannheim) and incubated at 37 °C in a humidified 5% CO2 atmosphere. Stable shMUS81 MRC5SV40 were described in Palma et al (21). The wild type form of RNAi-resistant MUS81 ORF cloned into the pEF1a-IRES-NEO plasmid was subjected to SDM (Quickchange II XL – Stratagene) to introduce the T86 or S97 mutations. Sequence-verified plasmids were then transfected into MRC5 shMUS81 cells by the Neon nucleofector (Life technologies). To generate shBRCA2 inducible cell lines, MRC5SV40 were transduced with lentivirus expressing an shBRCA2 cassette under the control of a Dox-regulated promoter at 0.5 of multiplicity of infection (MOI) (Dharmacon SmartVector inducible lentivirus, sequence code V3SH7669-225202141; (46)). After puromycin selection at 300 ng/ml, a single clone was selected and used throughout the study.

### Transfections

RNA interference against MUS81 was performed using FlexiTube HsMUS81 6 (Qiagen) at final concentration of 10 nM using Lullaby 48 h before to perform experiments. BRCA2 was depleted with ON-TARGETplus Human BRCA2-SMARTpool (Dharmacon) at 25 nM using Interferin (Polyplus) and experiments were performed after 48 h of transfection. RNA interference against EME2 was performed with SMARTPOOL siRNA della Dharmacon (geneID 197342). The efficiency of protein depletion was monitored by western blotting 72 h after transfection. MRC5, MRC5 shMUS81/MRC5 shMUS81 or shMUS81 S97A cell lines were transfected with pcDNA3-HA-Chk2 (wild-type or with CHK2 mutation) using Neon transfection system 48 h prior to perform experiments.

### Chemicals

HU was added to culture medium at 2 mM from stock solutions 200 mM prepared in Phosphate-buffer saline solution (PBS) to induce DNA replication arrest or DNA damage. MIRIN, the inhibitor of MRE11 exonuclease activity (Calbiochem), was added to DMEM 15 min prior HU at 50 µM. Nocodazole (Sigma-Aldrich) was used at 0.5 μg/μl. The BML-277 CHK2 inhibitor (Calbiochem) was used at 10 µM. CldU (Sigma-Aldrich) was dissolved in sterile water at 200mM stock solution and used at 50 μM. IdU (Sigma-Aldrich) was dissolved in sterile DMEM as a stock solution 2.5 mM and stored at −20 °C. RAD52 inhibitor, EGC (Sigma-Aldrich) was dissolved in DMSO at 100 mM, and stock solution was stored at - 80 °C and was used at 50 µM.

### Phosphorylation site prediction

For the prediction of MUS81 phosphorylation sites, the MUS81 protein sequence was scanned using the GPS 2.0 software (http://gps.biocuckoo.org/download.php) with a medium threshold that consist in a < 6% of false identification rate (60)

### Production and purification of GST-fused fragments

The FHA domain of CHK2 was amplified by PCR from the pcDNA3-HA-CHK2 plasmid containing the full-length wild-type or mutant CHK2 ORF (a gift from Dr. Domenico Delia’s lab). PCR-amplified DNA was cloned into the GST-pGEX-2TK plasmid (Stratagene). GST-FHA fragment was expressed into E.coli BL21PlysS at 30°C for 4h. Bacterial pellets were lysed in BER reagent (Pierce) supplemented with DNase and Lysozyme, as indicated by the manufacturer, and purified by incubation with GSH-magnetic beads (Promega) after extensive washings, as reported previously (61). Quantification of the magnetic-beads-bound fragments was performed after SDS-PAGE and Coomassie staining against serial dilutions of purified BSA. Magnetic beads-bound GST-FHA fragments were then used as substrates for pull-down assays.

### Western blot analysis

Blots were incubated with primary antibodies against: anti-MUS81 (Santa Cruz Biotechnology, 1:2000), anti-Lamin B1 (Abcam, 1:20000), anti-CHK2 (Santa Cruz Biotechnology, 1:1000), anti-BRCA2 (Bethyl, 1:2000), anti-pCHK2 (Millipore, 1:2000), anti-SMARCAL1 (Bethyl, 1:2000), anti-GFP (Santa Cruz Biotechnology, 1:1000), anti-pS1981ATM (Cell signaling, 1:1000), anti-ATM (Cell signaling, 1:1000), anti-HA (Cell signaling, 1:1000), anti-Cyclin A (Santa Cruz Biotechnology, 1:1000), anti-EME1 (Santa Cruz Biotechnology, 1:500), anti-EME2 (Thermo Fisher, 1:1000), anti-FLAG (Sigma Aldrich, 1:2000). After incubations with horseradish peroxidase-linked secondary antibodies (Jackson ImmunoResearch, 1:30,000), the blots were developed using the chemiluminescence detection kit ECL-Plus (Amersham) according to the manufacturer’s instructions. Quantification was performed on blot acquired by ChemiDoc XRS+ (Bio-Rad) using Image Lab software, and values shown on the graphs represent a normalisation of the protein content evaluated through Lamin B1 or GAPDH immunoblotting.

### Immunofluorescence

Cells were grown on 35-mm coverslips and harvested at the indicated times after treatments. For immunofluorescence (IF), after further washing with PBS, cells were pre-extracted with 0.5% Triton X-100 and fixed with 3% para-formaldehyde (PFA)/2% sucrose at room temperature (RT) for 10 min. After blocking in 3% bovine serum albumin (BSA) for 15 min, staining was performed with mouse monoclonal anti-phospho-ATM Ser1981 (Millipore 1:300) diluted in a 1% BSA/0.1% saponin in PBS solution, for 1 h at 37 °C in a humidifier chamber. After extensive washing with PBS, Alexa Fluor® 488 conjugated-goat anti-mouse was used at 1:200 and applied for 1 h at RT followed by counterstaining with 0.5 mg/ml 4,6-diamidino-2-phenylindole (DAPI).

### Detection of ssDNA by native IdU assay

To detect nascent ssDNA cells were labelled with 80 µM IdU for 20 min before treatments. Untreated cells were labelled for 20 min prior to performing immunofluorescence. For IdU detection cells were subjected to extensive washing with PBS, permeabilized with 0,5% TRITON-X 100 at 4° for 8 min and then fixed with 3% PFA/ 2% Sucrose. Fixed cells were blocked with 3% BSA for 20 min and incubated with primary antibody anti-IdU antibody (Becton Dickinson 1:100) for 1 h at 37 °C in 1% BSA/PBS in a humid chamber, followed by species-specific fluorescein-conjugated secondary antibody Alexa Fluor 488 (Goat Anti-Mouse IgG (H + L), highly cross-adsorbed—Life Technologies) diluted 1:200 in 1% BSA/PBS for 1 h at room temperature. After extensive washing with PBS, cells were stained with 0.5 mg/ml 4,6-diamidino-2-phenylindole (DAPI). Slides were analysed with Eclipse 80i Nikon Fluorescence Microscope, equipped with a Video Confocal (ViCo) system. For each time point, at least 100 nuclei were examined by two independent investigators and foci were scored at 40×. Quantification was carried out using the ImageJ software.

### In situ PLA assay for protein–protein interaction

The in-situ PLA (DuoLink, Merck) was performed according to the manufacturer’s instructions. After indicated treatments, cells were permeabilized with 0.5% Triton X-100 for 10 min at 4 °C, fixed with 3% PFA/2% Sucrose solution for 10 min and then blocked in 3% BSA/PBS for 15 min. After washing with PBS, cells were incubated with the two relevant primary antibodies. The primary antibodies used were as follows: mouse monoclonal anti-FLAG (SIGMA 1:100), rabbit polyclonal anti-CHK2 (Santa Cruz, 1:70). Samples were incubated with secondary antibodies conjugated with PLA probes MINUS and PLUS: the PLA probe anti-mouse PLUS and anti-rabbit MINUS. The incubation with all antibodies was accomplished in a humidified chamber for 1 h at 37 °C. Next, the PLA probes MINUS and PLUS were ligated using two connecting oligonucleotides to produce a template for rolling-circle amplification. After amplification, the products were hybridised with red fluorescence-labelled oligonucleotide. Samples were mounted in Prolong Gold anti-fade reagent with DAPI (blue). Images were acquired randomly using Eclipse 80i Nikon Fluorescence Microscope, equipped with a Video Confocal (ViCo) system.

### Quantitative in situ analysis of protein interactions at DNA replication forks (SIRF assay)

10.000 cells were plated onto chamber slides the day before the experiment. The day of experiment, cells were incubated with 125 µM EdU for 8 min, followed 2mM HU as indicated. After treatments, cells were fixed with 2% PFA/PBS for 15 min at RT. Cells were permeabilized in 0,25% Triton X-100 in PBS for 15 min at room temperature. Click reaction cocktail was added into slides (30 µl/well) and incubated at room temperature 30 min in a humid chamber protected from light. Slides were washed with PBS and blocked in 3% BSA/PBS for 20 min. After washing with PBS, cells were incubated with the primary antibodies. The primary antibodies used were rabbit polyclonal anti-Biotin (Abcam 1:500) and mouse monoclonal anti-phospho-ATM Ser1981 (Millipore 1:300). Samples were incubated with secondary antibodies conjugated with PLA probes MINUS and PLUS: the PLA probe anti-mouse PLUS and anti-rabbit MINUS (OLINK Bioscience). The incubation with all antibodies was accomplished in a humidified chamber for 1 h at 37 °C. Next, the PLA probes MINUS and PLUS were ligated using two connecting oligonucleotides to produce a template for rolling-circle amplification. After amplification, the products were hybridised with red fluorescence-labelled oligonucleotide. Samples were mounted in Prolong Gold anti-fade reagent with DAPI (blue). Images were acquired randomly using Eclipse 80i Nikon Fluorescence Microscope, equipped with a Video Confocal (ViCo) system.

### DNA Fiber assay

Cells were pulse-labelled with 50 µM CldU for 20 min, washed twice with PBS and treated with HU 2mM as reported in the experimental schemes. After treatments cells were washed twice with PBS and incubated for 40 min in fresh medium with 250 μM IdU. DNA fibres were prepared and analysed to evaluate fork restart (40). For immunodetection of labelled tracks, the following primary antibodies were used: rat anti-CldU/BrdU (Abcam) and mouse anti-IdU/BrdU (Becton Dickinson).

### Chromatin isolation

Cells (4 × 10 × 6 cells/ml) were resuspended in buffer A (10 mM HEPES, [pH 7.9], 10 mM KCl, 1.5 mM MgCl2, 0.34 M sucrose, 10% glycerol, 1 mM DTT, 50 mM sodium fluoride, protease inhibitors [Roche]). Triton X-100 (0.1%) was added, and the cells were incubated for 5 min on ice. Nuclei were collected in pellet by low-speed centrifugation (4 min, 1300 × *g*, 4 °C) and washed once in buffer A. Nuclei were then lysed in buffer B (3 mM EDTA, 0.2 mM EGTA, 1 mM DTT, protease inhibitors). Insoluble chromatin was collected by centrifugation (4 min, 1700 × *g*, 4 °C), washed once in buffer B + 50 mM NaCl, and centrifuged again under the same conditions. The final chromatin pellet was resuspended in 2X Laemmli buffer and sonicated for 15 s in a Tekmar CV26 sonicator using a microtip at 25% amplitude.

### Neutral comet assay

DNA breakage induction was evaluated by Comet assay (single-cell gel electrophoresis) in non-denaturing conditions. Briefly, dust-free frosted-end microscope slides were kept in methanol overnight to remove fatty residues. Slides were then dipped into molten low melting point (LMP) agarose at 0.5% and left to dry. Cell pellets were resuspended in PBS and kept on ice to inhibit DNA repair. Cell suspensions were rapidly mixed with LMP agarose at 0.5% kept at 37 °C and an aliquot was pipetted onto agarose-covered surface of the slide. Agarose-embedded cells were lysed by submerging slides in lysis solution (30 mM EDTA, 0.1% sodium dodecyl sulphate (SDS)) and incubated at 4 °C, 1 h in the dark. After lysis, slides were washed in Tris Borate EDTA (TBE) 1X running buffer (Tris 90 mM; boric acid 90 mM; EDTA 4 mM) for 1 min. Electrophoresis was performed for 20 min in TBE 1X buffer at 1 V/cm. Slides were subsequently washed in distilled H_2_O and finally dehydrated in ice-cold methanol. Nuclei were stained with GelRed (1:1000) and visualised with a fluorescence microscope (Zeiss), using a 60X objective, connected to a CCD camera for image acquisition. At least 300 comets per cell line were analysed using CometAssay IV software (Perceptive instruments) and data from tail moments processed using Prism software. Apoptotic cells (smaller Comet head and extremely larger Comet tail) were excluded from the analysis to avoid artificial enhancement of the tail moment.

