## Supplementary Figures and Legends for "CHK2 regulates MUS81-dependent DSBs in response to replication stress and BRCA2 deficiency"

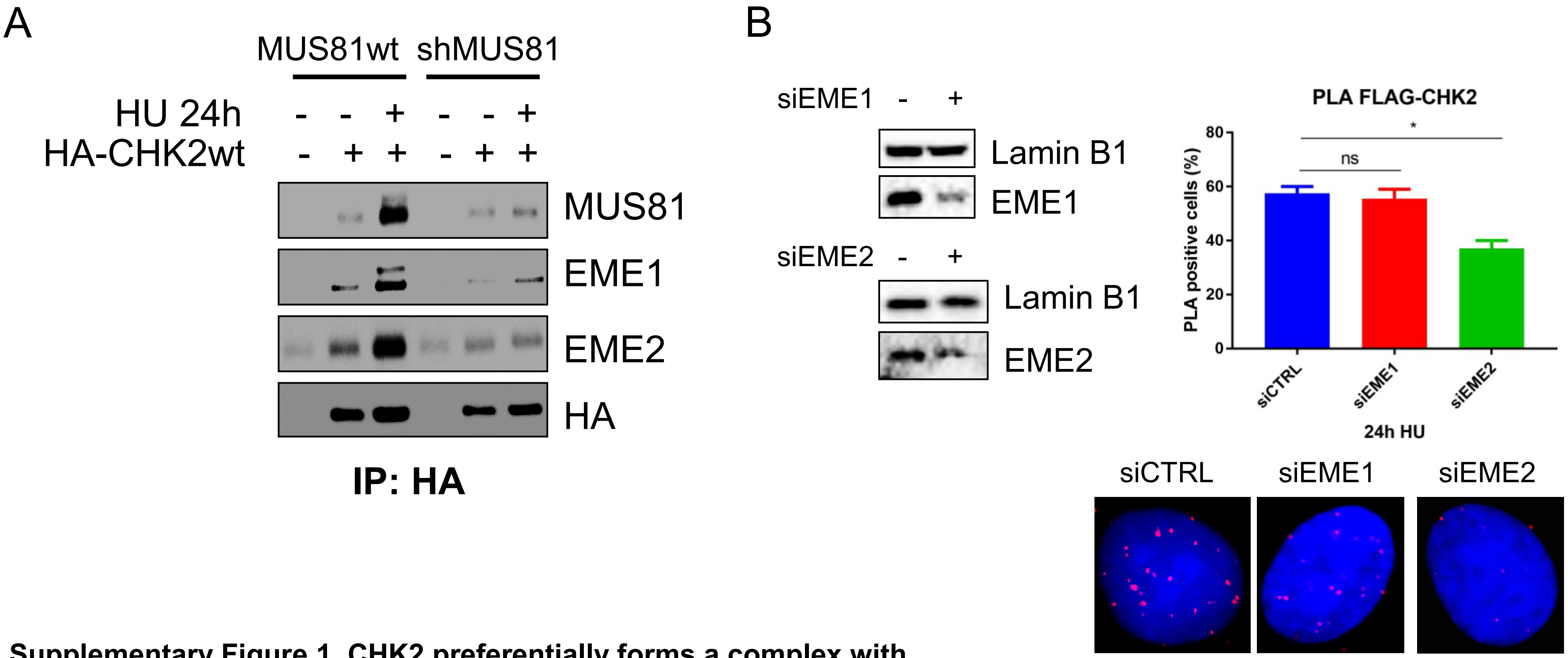


**Supplementary Figure 1. CHK2 preferentially forms a complex with the MUS81-EME2 heterodimer after replication stress.** A) MRC5 shMUS81 cells stably complemented with Flag-MUS81 were transfected with the HA-CHK2-expressing plasmid or empty plasmid and used to immunoprecipitate CHK2 with anti-HA antibodies in cells treated with HU as indicated. B) MRC5 shMUS81 cells stably complemented with Flag-MUS81 were depleted of EME1 or EME2 by transient transfection with siRNA oligos as indicated and the interaction between Flag-tagged MUS81 and CHK2 evaluated by *in situ* PLA after treatment with HU 2mM using the anti-FLAG and anti-CHK2 pair. The blots show the depletion level of each MUS81 partner. Images of representative single nuclei from each condition are shown. The graph shows the quantification of PLA-positive cells for each genotype (mean±SE; n=2. * = p < 0,5 ANOVA).

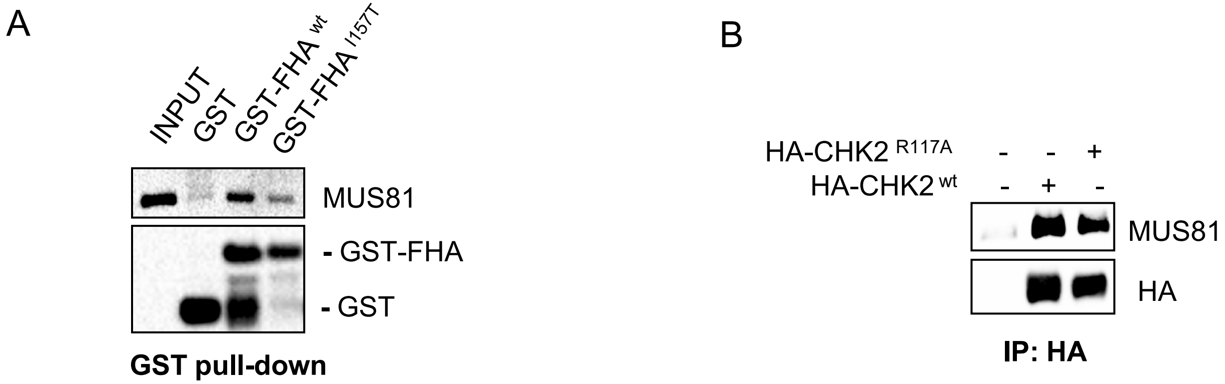


**Supplementary Figure 2. Interaction of MUS81 and CHK2 FHA requires I157.** A) The wild-type and I157T-mutated GST-fused FHA fragment of CHK2 was purified from bacteria and used as bait to pull-down MUS81 from HEK293T nuclear extracts. B) MRC5 shMUS81 cells stably complemented or not with Flag-MUS81 were transiently transfected with the indicated HA-CHK2 construct and used to immunoprecipitate CHK2 in cells treated with HU. Representative immunoblots are shown.

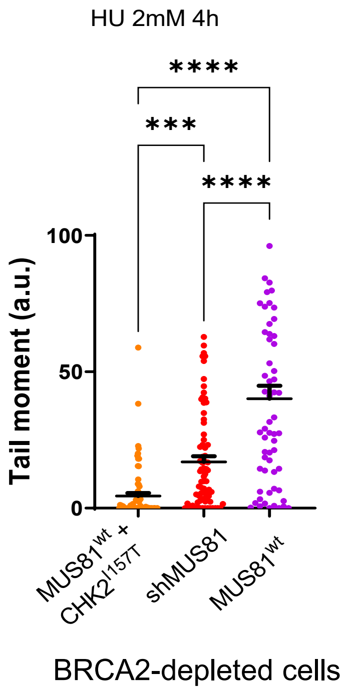


**Supplementary Figure 3. Depletion of MUS81, impaired interaction of MUS81 and CHK2 or inhibition of ATM similarly reduces formation of DSBs induced by BRCA2-depletion.** MRC5 shMUS81 cells stably complemented with Flag-MUS81 were transfected with BRCA2 siRNA alone or in combination with the mutant HA-CHK2-expressing plasmid and treated with HU for 4h. Formation of DSBs was analysed by neutral Comet assay. (ns = not significant; *** = $p<0.01$; **** = $p<0.001$ Student's t-test).

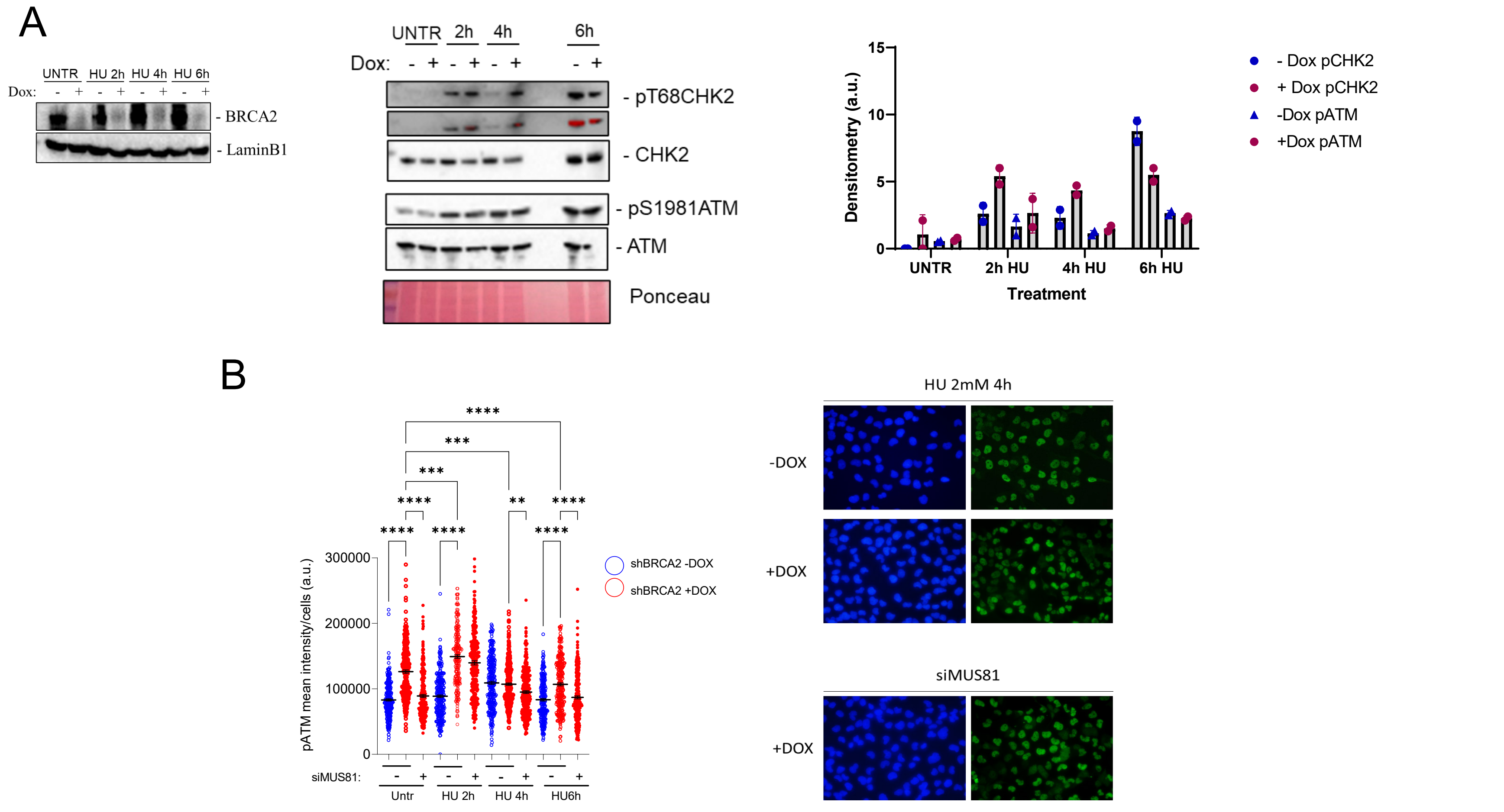


**Supplementary Figure 4. The ATM-CHK2 axis is activated early in the absence of BRCA2 independently on MUS81.** A) Western blot analysis of BRCA2 in MRC5 shBRCA2 inducible cell lines in presence or not of doxycycline. Lamin B1 was used as a loading control. Western blot analysis of ATM and CHK2 activation in total extracts of MRC5 shBRCA2 ± doxycycline. Phosphorylation of ATM and CHK2 was assessed using anti-pATM (S1981), and anti-pCHK2 (T68) antibody. Total amount of ATM, and CHK2 was determined with anti-ATM or anti-CHK2 antibody. Ponceau was used to normalize for total protein. B) Evaluation of ATM activation by immunofluorescence in MRC5 shBRCA2 cells line after transfection of siMUS81 oligos. The presence of activated ATM was assessed using S1981 phospho-specific antibody pATM (S1981). Nuclei were counterstained with DAPI. Representative images of cells stained for pATM and DAPI are given. Dot plot shows pATM intensity per nucleus from three independent experiments (**** = $P < 0.0001$; *** = $P < 0.001$; two-tailed Student's t test).

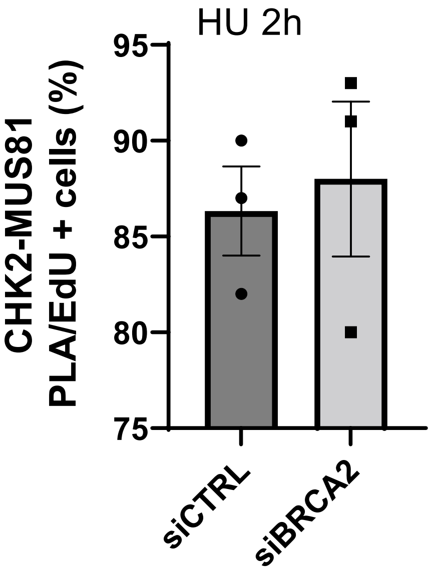


**Supplementary Figure 5. Interaction between CHK2 and MUS81 mostly occurs in S-phase in HU-treated cells.** MRC5 shMUS81 cells stably complemented with Flag-MUS81 were transfected with BRCA2 siRNA alone or in combination with the wild-type HA-CHK2-expressing plasmid and treated with HU. Immediately before treatment, EdU was added to cells to label S-phase. Formation of CHK2-MUS81 complexes was analysed by *in situ* PLA followed by Click-It-mediated detection of EdU. The graph shows the number of PLA-positive cells showing at least 5 spots that are also EdU-positive. Values are means ±S.E; n=2.

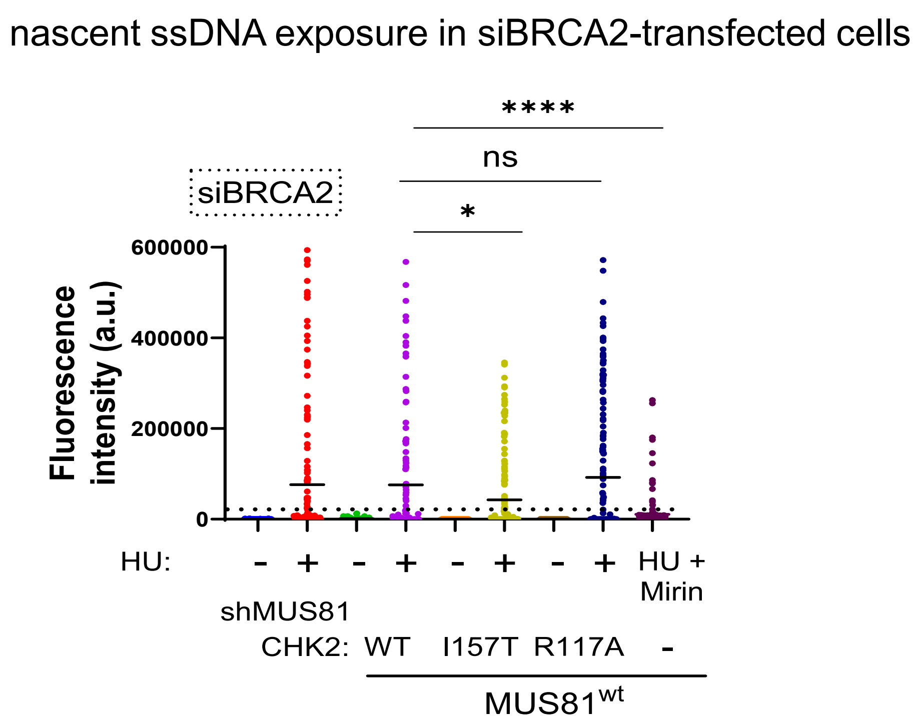


**Supplementary Figure 6. MRE11-dependent exposure of nascent ssDNA in the absence of BRCA2 is not affected by loss of MUS81-CHK2 function.** MRC5 shMUS81 cells stably complemented with Flag-MUS81 were transfected with BRCA2 siRNA alone or in combination with a plasmid expressing the wild-type or FHA mutant form of CHK2. After 48h, cells were treated with 2mM HU for 4h. As a control, cells were co-treated with Mirin. Immediately before treatment, IdU was added to cells to label nascent DNA. Formation of ssDNA was analysed by native anti-IdU immunofluorescence. The dotted line shows the mean level of nascent ssDNA detected in BRCA2-proficient cells. The graph shows the quantification of ssDNA intensity. Mean values ±S.E. are indicated. (n=2; ns= not significant; *=p<0.5; **** =p<0.001, ANOVA).

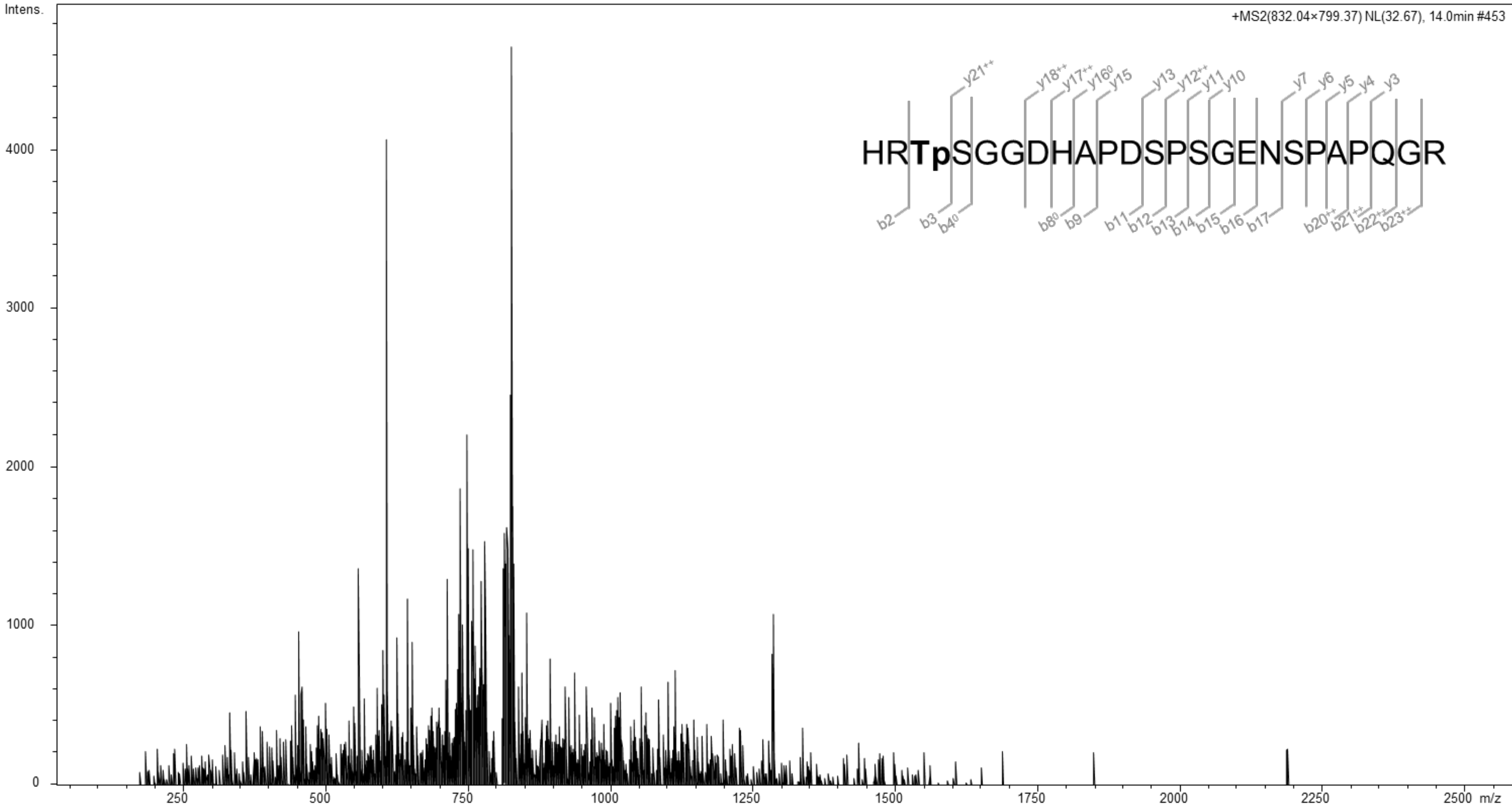


**Supplementary Figure 7.** CID MS/MS spectrum of the triply charged parent ion of the tryptic phosphopeptide HRTSGGDHAPDSPSGENSPAPQGR (m/z 832.04) from MUS81 protein. The observed b-and y-type fragment ions are shown with the peptide sequence in the insert. Tp corresponds to phosphorylated Thr-86.

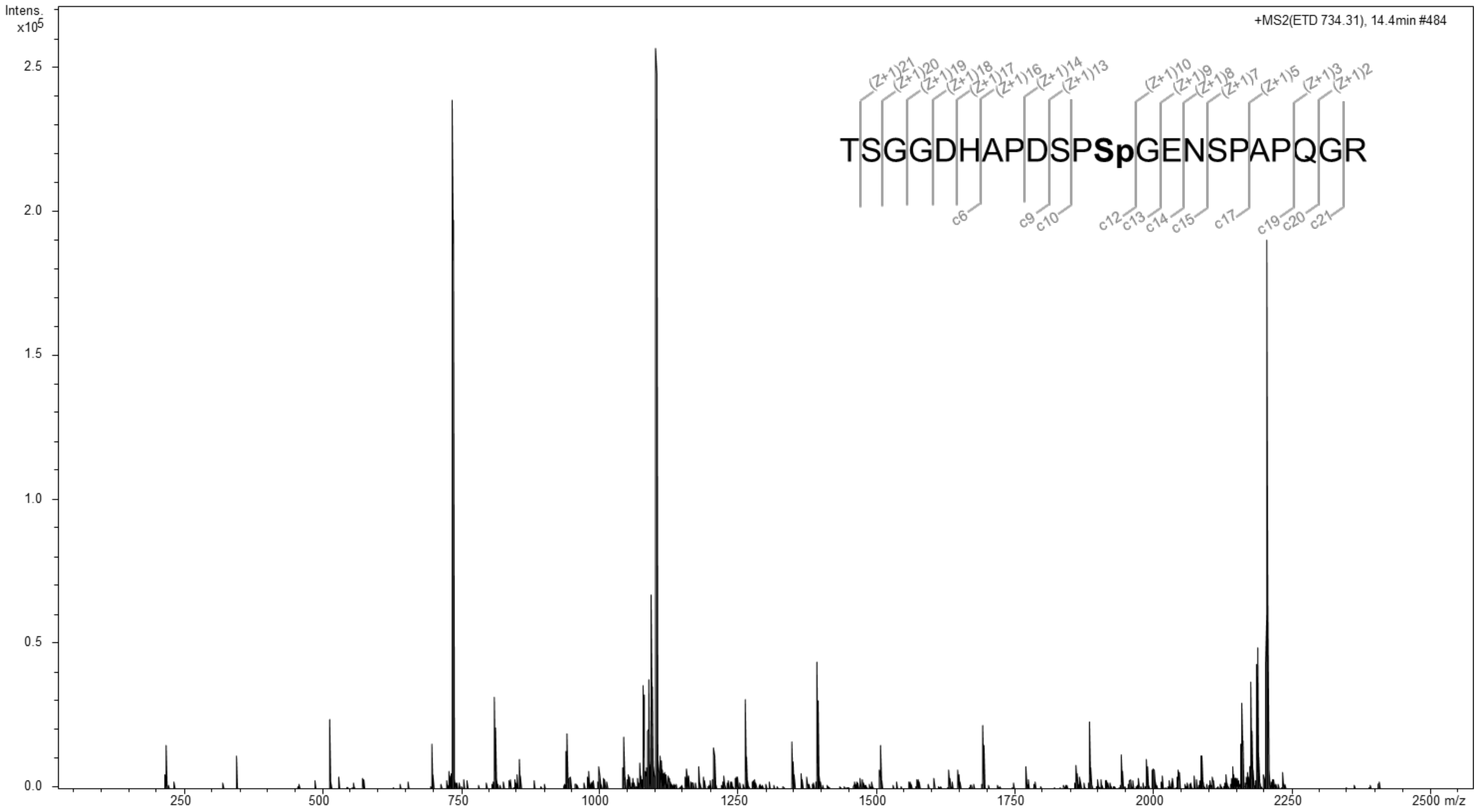


**Supplementary Figure 8.** ETD MS/MS spectrum of the triply charged parent ion of the tryptic phosphopeptide TSGGDHAPDSPSGENSPAPQGR (m/z 734.31) from MUS81 protein. The observed c-and z-type fragment ions are shown with the peptide sequence in the insert. Sp corresponds to phosphorylated Ser-97.
